# Ratiometric RNA labeling allows dynamic multiplexed analysis of gene circuits in single cells

**DOI:** 10.1101/2021.09.23.461487

**Authors:** Shuhui Xu, Kai Li, Liang Ma, Jianhan Zhang, Shinae Yoon, Michael B. Elowitz, Yihan Lin

## Abstract

Biological processes are highly dynamic and are regulated by genes that connect with one and another, forming regulatory circuits and networks. Understanding how gene regulatory circuits operate dynamically requires monitoring the expression of multiple genes in the same cell. However, it is limited by the relatively few distinguishable fluorescent proteins. Here, we developed a multiplexed real-time transcriptional imaging method based on two RNA stem-loop binding proteins, and employed it to analyze the temporal dynamics of synthetic gene circuits. By incorporating different ratios of MS2 and PP7 stem-loops, we were able to monitor the real-time nascent transcriptional activities of up to five genes in the same cell using only two fluorescent proteins. Applying this multiplexing capability to synthetic linear or branched gene regulatory cascades revealed that propagation of transcriptional dynamics is enhanced by non-stationary dynamics and is dictated by the slowest regulatory branch in the presence of combinatorial regulation. Mathematical modeling provided further insight into temporal multi-gene interactions and helped to understand potential challenges in regulatory inference using snapshot single-cell data. Ratiometric multiplexing should scale exponentially with additional labelling channels, providing a way to track the dynamics of larger circuits.

## INTRODUCTION

Gene regulatory networks play critical roles in orchestrating diverse biological processes^1,2^, from embryo development to tissue homeostasis, and network malfunction can lead to abnormal phenotypes and diseases^3–5^. To interfere with dysregulated processes and to precisely engineer cells, it is critical to understand the structure and function of the underlying gene regulatory networks or the core circuits in the networks^6–10^. Complementary to population-level assays that investigate networks by averaging across cells, single-cell approaches can offer unique insights into the structure and function of gene circuits and networks^11–17^. Nevertheless, new single-cell approaches, especially those that provide temporal dynamical information, are needed to systematically understand circuits and networks.

A growing list of studies has demonstrated the crucial roles of temporal dynamics in gene circuits and networks. For example, it has been shown that some critical biological processes are controlled by regulatory circuits that are highly dynamic, including differentiation^18^, development^19^, and stress response^20^. In these systems, the temporal dynamics of regulators, rather than the levels of their expression, have been suggested to control cell fate^11–13^. More recently, coordinated transcriptional bursting of multiple genes within a network has been suggested to underlie the transition into physiologically important cell states such as drug resistance in cancer^21,22^. Thus, studying the temporal dynamics of circuits and networks at the single-cell level is necessary for understanding their functions.

Arguably the most direct approach for analyzing gene circuit dynamics would be to monitor the temporal changes in gene expression for all circuit components within the same single cell^23–25^. Along this direction, multiplexed time-lapse imaging techniques based on fluorescent reporter proteins have been developed for real-time quantifications of multiple genes of interest in natural or synthetic circuits^26–30^. However, this type of technique faces limitations, including the limited number of distinguishable fluorescent proteins^31^ and the difficulty of capturing fast gene regulatory signals. The latter limitation arises because fluorescent protein expression level reflects both transcriptional and post-transcriptional regulation, and fast transcriptional dynamics could be averaged away. Furthermore, the relatively slow and variable maturation rates of fluorescent proteins add an additional layer of complexity^23,32,33^. Therefore, alternative methodologies for monitoring fast temporal dynamics of gene circuits would be desirable.

In this work, towards the goal of temporal and scalable interrogation of gene circuits, we introduce a multiplexed transcriptional reporter system based on two phage-derived RNA binding proteins, MCP (bacteriophage MS2 coat protein^34,35^) and PCP (bacteriophage PP7 coat protein^36^). These two RNA binding proteins have been used in conjunction with their cognate RNA stem-loops to enable real-time quantitative imaging of transcriptional activities in different biological systems across organisms^34,35,37,38,36,39–44^. Based on the orthogonality of these two RNA binding protein systems, which was demonstrated in previous work^39,45–49^, we devised a ratiometric barcoding scheme by incorporating different ratios of MS2 and PP7 stem-loops, and achieved multiplexed reporting of the transcriptional activities of up to five genes in a circuit using only two fluorescent colors. This methodology allowed us to simultaneously monitor multi-gene interactions over a long period of time inside the same single mammalian cells. Through temporal, multiplexed analysis of synthetic gene circuits containing linear or branched regulatory cascade, we uncovered characteristics and limits for transcriptional burst propagation, and discovered that burst propagation is imbalanced for different branches of a cascade. The quantitative time-lapse data also allowed us to extract kinetic parameters, and to construct mathematical models that recapitulate the measured dynamics. Together, we envision that this method, as well as the integrated approach for analyzing gene circuit dynamics presented here, should enable multiplexed temporal decoding of other gene regulatory systems in single cells.

## RESULTS

### Design of a multiplexed transcriptional reporter system

To analyze the fast temporal dynamics in gene circuits using time-lapse imaging, we need a reporter system that (a) can report real-time transcriptional activities of genes; (b) is scalable to allow multiplexed temporal analysis of multiple genes in the same cell; and (c) uses as few fluorescent channels as possible in order to reduce phototoxicity caused by light illumination.

After surveying existing live-cell transcriptional reporter systems^50,51^, we chose to construct our system based on two widely-used RNA binding protein systems MCP/MS2 and PCP/PP7, because they have been shown to enable co-localized and quantitative detection of single mRNAs^39^. To enable multiplexing, we leveraged the spatial dimension, as the nascent transcription occurs in a defined genomic locus, and proposed a ratiometric barcoding scheme (**Fig. 1A**) that would allow quantifying multiple spatially-separated nascent transcription sites (**Fig. 1B**). The fluorescence intensity ratios of these nascent sites would be uniquely mapped to individual genes in a circuit, enabling temporal, multiplexed circuit analysis (**Fig. 1C**). Such a reporter system would satisfy the preceding requirements as follows: a) RNA binding protein-based transcriptional reporter systems allow quantifications of nascent transcriptional activities; b) these systems are modular and can be used in combination, thus enabling scalability and multiplexability; c) only two fluorescent proteins that are separately fused to MCP and PCP are needed.

**Figure 1.**
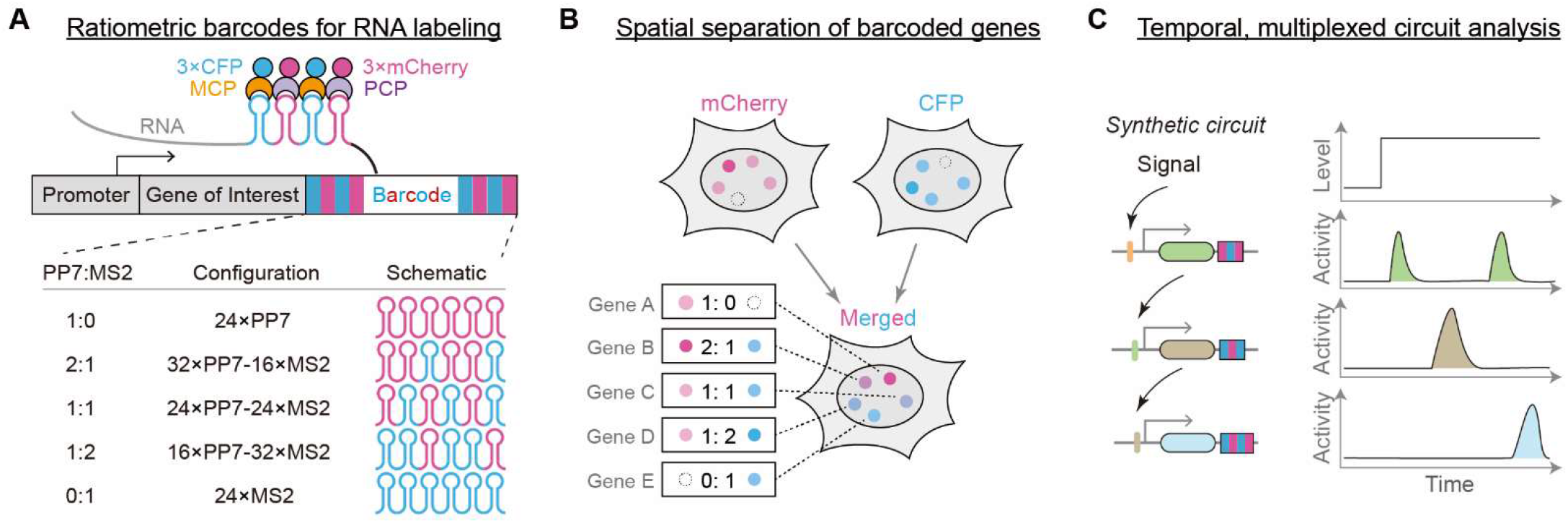
Conceptual scheme: MS2 and PP7 allow live cell dynamic transcriptional readouts of multiple genes. **(A-B)** Design of the reporter system. Gene of interest is fused with a barcode cassette composed of two types of RNA stem-loops, PP7 and MS2, which would recruit cognate binding proteins (i.e., PCP-3xmCherry and MCP-3xCFP) when transcribed (A, top). The ratio between the numbers of PP7 and MS2 stem-loops provides a ratiometric barcoding mechanism (A, bottom). Because genes with different barcodes are integrated into distinct chromatin loci, the spatial separation between these loci allows reading out the barcode identity as well as the transcriptional activity of each gene locus using two fluorescent channels (B). **(C)** Multiplexed temporal analysis of a synthetic circuit (schematic). The ratiometric barcodes would allow reading out temporal transcriptional activities of all genes in the circuit.

### Testing the multiplexed transcriptional reporter system

As a pilot experiment, we transiently transfected reporter plasmids into U2OS cells and tested whether different barcodes could be distinguished (**Methods**; see **Supplementary Fig. 1A-B** for illustration of experimental and analysis workflows). In cells separately transfected with each of five barcode-containing genes, we observed clear nascent transcription sites upon the addition of the inducer (doxycycline), where barcodes were bound by MCP and/or PCP molecules (**Supplementary Fig. 1C**). The two RNA binding proteins are indeed orthogonal in our cell line as previously reported ^39,45–49^, i.e., PCP does not bind to MS2 arrays while MCP does not bind to PP7 arrays (**Supplementary Fig. 1C**). Further quantifications of the two fluorescent protein intensities at the nascent transcription sites showed clear separations of the five different barcodes (**Supplementary Fig. 1D**).

Because temporal circuit analysis requires time-lapse imaging for a long duration (> 10 hours), we next constructed stable cell lines containing ratiometrically barcoded reporter genes and tested the long-term performance of the barcodes using microscopy (see **Supplementary Fig. 1A** for workflow). The integration of barcoded genes was accomplished by using piggyBac transposase (**Methods**). Reassuringly, the RNA stem-loop cassettes provided multiplexing capabilities in stable cell lines (**Fig. 2A-B, Supplementary Fig. 2A**) as in transiently transfected cells (**Supplementary Fig. 1C-D**), and we could capture the temporal activation and deactivation events at the nascent transcription sites (**Fig. 2A**, **Supplementary Video 1**).

**Figure 2.**
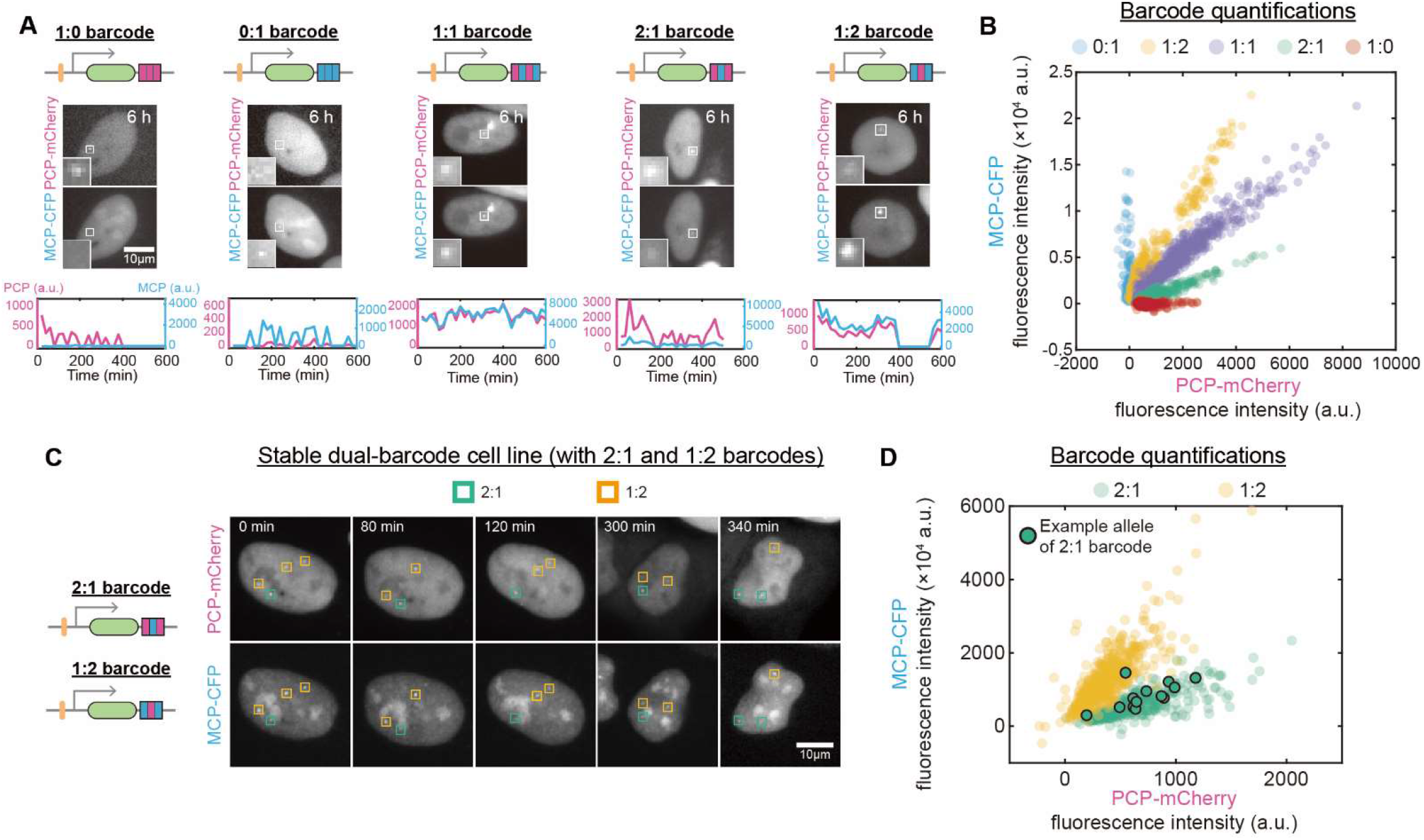
Five-channel transcriptional imaging using two fluorescent protein colors. **(A-B)** Characterizations of cells stably integrated with individual barcode-labeled genes. Snapshots and temporal fluorescence intensity traces of representative single cells stably integrated with genes fused with indicated barcodes are shown (A). Fluorescence intensities of all barcodes were plotted together in (B) (n = 19, 20, 23, 20 and 22 cells respectively for 1:0, 2:1, 1:1, 1:2 and 0:1 barcodes). Doxycycline (1 μg/mL) was added before t = 0 min. See also **Supplementary Fig. S2A** and **Supplementary Video 1**. **(C-D)** Transcriptional imaging of two barcodes in the same cell. Two-color time-lapse images of a representative single cell integrated with two different genes carrying distinct barcodes (2:1 and 1:2 barcodes) (C), and the fluorescence intensities of barcoded gene loci (D, n = 14 cells) are shown. The highlighted data points in (D) were from the temporal intensity traces of one example 2:1 barcoded gene, showing that barcode identity needs to be determined using data from multiple time points (as data from specific time points may deviate from the averaged behavior). Doxycycline (1 μg/mL) was added before t = 0 min. See also **Supplementary Fig. S2B-C** and **Supplementary Video 2**.

We further asked whether multiple ratiometric barcodes could be resolved inside the same single cells during imaging. We constructed stable cell lines, each integrated with two genes separately fused with distinct barcodes. By conducting time-lapse imaging on these cell lines, we could monitor and track the nascent transcription sites of different genes in the same cells over time (**Fig. 2C**, **Supplementary Fig. 2B**, **Supplementary Video 2**), and resolve spatially separated barcodes with two fluorescent colors (**Fig. 2D**, **Supplementary Fig. 2C**). Note that we often observed multiple integration events for each barcoded gene (e.g., **Supplementary Fig. 2B**), a commonly observed outcome for piggyBac-mediated plasmid integration^52^. Furthermore, temporal recordings of the barcodes are necessary for the accurate determination of barcode identities, as data from single time points might deviate from the average behavior (see **Fig. 2D** for an example allele). Together, these data indicated that we could use the barcodes to analyze the temporal dynamics of multi-gene circuits (**Fig. 1C**).

### Multiplexed analysis of a synthetic three-gene circuit

We sought to study the temporal behaviors of gene circuits with the reporter system. We focused on synthetic gene circuits, as we could analyze synthetic circuits with known regulatory interactions and relatively low interference from the host cell, allowing us to extract quantitative parameters and construct mathematical models of circuit dynamics. Additionally, because static single-cell expression data has been widely used for inferring regulatory relationships between genes^17,53^, temporal data from synthetic circuits would allow us to analyze whether and how the measurement timing of the snapshot data affects the inference of regulatory interactions, which could help to identify limitations when reconstructing regulatory relationships using static data.

Because two prominent features of human gene regulatory network are the hierarchical organization of transcription factors and the combinatorial regulation of genes^54^, we sought to construct and analyze synthetic circuits that recapitulate these features. We first constructed a monoclonal U2OS cell line containing a three-gene circuit constituting a hierarchical transcription factor cascade, where the external signal (doxycycline) is transmitted to an ‘effector’ protein (i.e., YFP) through two synthetic regulators, LacI-VP64 and Gal4-VP64 (**Fig. 3A**). These synthetic regulators were constructed with microbial (LacI, *lac* repressor from E. coli; Gal4, DNA binding domain of Gal4 from budding yeast) or viral components (VP64, four tandem copies of VP16 from Herpes simplex virus). For simplicity, we hereafter refer to the first regulator LacI-VP64 as gene A, the second regulator Gal4-VP64 as gene B, and the third protein YFP as gene C. For multiplexed detection, genes were fused with distinct ratiometric barcodes that can be distinguished by MCP and PCP intensity ratios (**Supplementary Fig. 3A**). Upon doxycycline addition, we observed that the three genes were transcriptionally activated in apparently sequential order over a time course of ~24 hours and the YFP signals emerged at later time points (**Fig. 3B-C**, **Supplementary Video 3**). Both population-averaged dynamics (**Supplementary Fig. 3B**) and cross-correlation functions (**Supplementary Fig. 3C**) indicated temporal signal transmission from gene A to gene B, and then to gene C, suggesting that this circuit transmits dynamic signals through cascading activation of circuit components.

**Figure 3.**
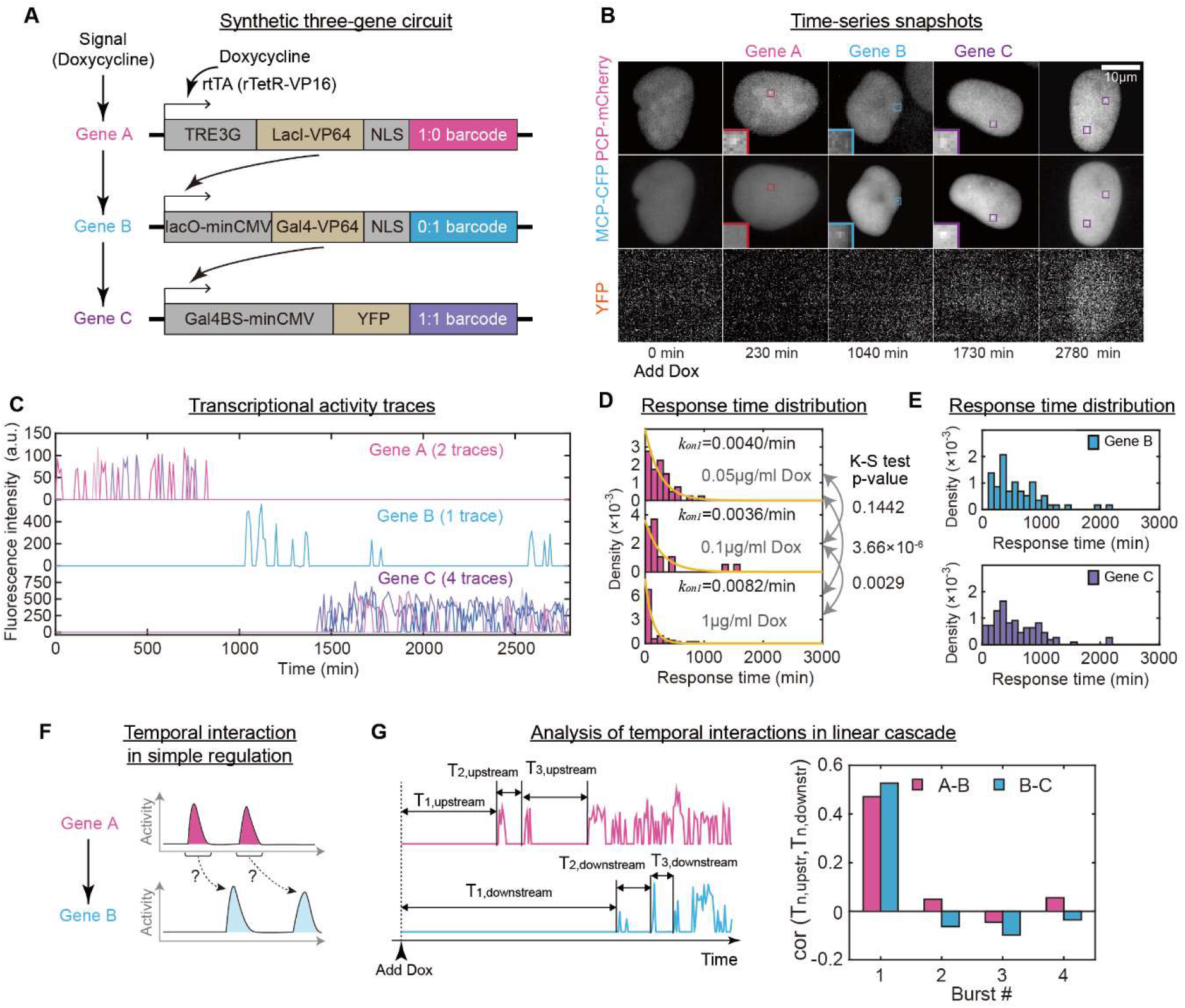
Analyzing dynamic multi-gene interactions in a synthetic three-gene circuit. **(A)** Design of a synthetic three-gene circuit. In this circuit, signal propagates from extracellular doxycycline to the first gene, LacI-VP64, and then to the second gene, Gal4-VP64, and lastly to YFP. Each gene is fused with a distinct barcode. **(B-C)** Time-lapse images (B) and corresponding transcriptional activity traces (C) of a representative single cell. Doxycycline (1 μg/ml) was added at t = 0 min. Note that traces of all integrated loci were plotted together. See also **Supplementary Video 3**. **(D-E)** Characterizations of the response time distributions. Response time distributions of gene A at different doxycycline concentrations are exponential and the estimated decay constant *k_on1_* represents the rate constant for the activation of the initial burst of gene A (D). n = 40, 19, 53 events for low, medium, high doxycycline, respectively. In contrast, the distributions for the other two genes are peaked (E). n = 58 (gene B) and 110 (gene C) events. **(F)** Illustration of temporal gene-gene interaction at the level of individual transcriptional bursts. See also **Supplementary Fig. 4A**. **(G)** Characterization of transcriptional burst propagation. When each burst of the upstream gene is perfectly propagated to its downstream gene, the burst intervals for both genes would be positively correlated. We thus used the correlation between bust intervals for both genes to quantify burst propagation. The burst interval for the n^th^ burst (Tn) is calculated as the time difference between the start of the preceding burst (or dox addition) to the start of the n^th^ burst (left). The correlations between burst intervals for two gene pairs (i.e., A/B and B/C) are shown for the four initial bursts (right). See also **Supplementary Fig. 4B**.

The data above prompted questions regarding the quantitative behaviors of the circuit dynamics, for example, how external signal temporally transmits along the cascade. To address this, we first focused on gene A of the cascade, which is activated by a regulator (i.e., rtTA^55^, rTetR-VP48) that is post-translationally switched on by the external signal, doxycycline. These regulatory interactions resemble how nuclear receptors activate downstream target genes in response to extracellular signals such as steroid hormones^56^.

By analyzing the time it takes for gene A to activate in response to the external signal (i.e., doxycycline addition), we found that the response time (i.e., the time from signal addition to the first transcriptional burst) of gene A is exponentially distributed (**Fig. 3D**), indicative of a rate-limiting reaction step dictating the transmission from doxycycline to gene A. Because doxycycline switches on the DNA binding capacity of the regulator rtTA and its concentration affects the response time of gene A (**Fig. 3D**, **Supplementary Fig. 3B**), this rate-limiting step could result from the rtTA-dependent alternation of the chromatin state. To test this, we performed the experiment in the presence of TSA (trichostatin A, a histone deacetylase inhibitor), and found that the response time distribution was perturbed (**Supplementary Fig. 3D**). In addition to delaying the response of gene A, TSA also reduced its expression level (compare **Supplementary Fig. 3E** with **Supplementary Fig. 3B**). These results suggest that chromatin state alteration could be the rate-limiting step for the activation of the gene from the initially off state. Such TF-dependent chromatin state alternation has been described in models of quantitative gene regulation^57,58^. We noted that TSA slowed down gene activation and reduced gene activation level, suggesting that inhibiting histone deacetylation could reduce the expression of certain genes^59–61^. Thus, these results indicate that epigenetic mechanisms likely play a key role in shaping the kinetics of target gene activation in this synthetic cascade.

### Analyzing temporal multi-gene interactions in the linear cascade

Because this synthetic circuit captures the hierarchical organization of human transcription factors^54^, understanding how the synthetic regulators temporally interact with each other could shed light on the temporal organization of human transcription factor network. To dissect the temporal interactions between synthetic regulators, we analyzed the propagation of transcriptional bursts in the cascade. Analogous to analyzing spike propagation between neurons in a neural circuit^62^, analyzing the propagation of bursts between genes in a gene circuit could reveal the extents and characteristics of temporal gene-gene interactions.

We first asked whether and how the initial transcriptional burst of gene A could propagate to gene B. The distribution of time delay between the first burst of gene A and the first burst of gene B (i.e., the response time distribution of gene B) was peaked (**Fig. 3E** top), as with the response time of gene C (**Fig. 3E** bottom). In contrast to the exponential response time distribution of gene A (**Fig. 3D**), peaked distributions indicate the presence of better-defined timescale for the transmission of dynamic signals, which is mostly unaffected by doxycycline concentration (**Supplementary Fig. 3F**). We reasoned that how upstream regulator is activated would affect the shape of response time distribution of the downstream gene. More specifically, the regulator for gene A (i.e., rtTA, **Fig. 3A**) was already expressed when doxycycline was added, and thus the regulator can quickly activate gene A without delay, whereas the regulators for B and C were induced to express and needed to accumulate to sufficient concentrations, resulting in a relatively well-defined time delay from the regulator’s first transcriptional burst to the target gene’s first transcriptional burst.

Importantly, the observation of peaked response time distribution may not necessarily indicate the presence of regulatory interactions, as such a distribution could also result from other scenarios, for example, two unrelated genes being independently and separately activated by two temporally spaced signals. Therefore, we reasoned that an alternative quantification of burst propagation is necessary for studying temporal gene-gene interaction at the level of individual bursts (**Fig. 3F**). We noticed that the relationship between burst starting times of two genes could allow assessing the presence or absence of gene-gene interaction (**Supplementary Fig. 4A**). We thus generated scatter plots of the starting time of the upstream gene’s first burst versus the starting time of its immediate downstream gene’s first burst, and found that for both gene pairs, the two starting times are significantly correlated (**Supplementary Fig. 4B**). The data points are along a line that is mostly parallel to the diagonal (**Supplementary Fig. 4B**), recapitulating the peaked response time distributions (**Fig. 3E**). These results indicate that the first burst of the upstream regulator gene can propagate to the downstream target gene.

To examine the propagation of subsequent bursts, we computed analogous correlations for additional bursts of upstream and downstream genes (**Fig. 3G** left). Unexpectedly, we found that the correlations are insignificant (**Fig. 3G** right), suggesting that these successive bursts of upstream genes were not effectively propagated downstream.

### Constructing a kinetic model using multiplexed temporal data

To explain these perplexing dynamics and gain further insight into the temporal circuit dynamics, we next sought to describe the circuit behaviors with a chemical kinetic model. We needed to use a three-state model to describe the transcriptional bursting dynamics of gene A (**Supplementary Fig. 5A**), whose state transition rate constants can be inferred from dwell time distributions (**Supplementary Fig. 5B**). The rationale for using a three-state model is because that the initial OFF state is different from the later OFF state based on the differences in their dwell times. More specifically, by examining the time from doxycycline addition to the first burst of gene A, we could infer the activation rate constant *k_on1_* for this reaction (**Fig. 3D**). By plotting the dwell time distributions for the subsequent burst events, we found that the distributions for both ON and OFF states are exponential (**Supplementary Fig. 5C**), indicative of the presence of single rate-limiting steps for both state transition reactions. Notably, the rate of OFF to ON switch (**Supplementary Fig. 5C** bottom) is much faster compared to the switch from the initial OFF state to ON (**Fig. 3D**), supporting the use of a three-state model. Mechanistically, the difference in the switching rates from the two OFF states to the ON state could arise from the difference in chromatin states, because cells were maintained in a doxycycline-free medium prior to experiments and the promoter had remained silenced for an extended period of time before adding doxycycline.

We next asked whether a three-state model is necessary for describing the dynamics of genes B and C, or in other words, whether all genes in the circuit need to be described by two OFF states. We thus needed to extract the rate constants for OFF to ON transitions for both genes B and C. As discussed above, the activation of the first transcriptional burst for gene A is kinetically different from that for gene B or C, as reflected by the response time distributions (**Fig. 3D-E**). For both genes B and C, their regulator concentrations are changing during the activation of their first bursts. Because these reactions are non-stationary, we performed maximum-likelihood estimations of their rate constants (**Supplementary Fig. 5D** and Methods). Of note, there appear to be two distinct types of gene B loci that have different activation kinetics (**Supplementary Fig. 5D** left, see Methods). For the subsequent bursts, we estimated the rate constants from the dwell time distributions (**Supplementary Fig. 5E**). By doing so, we found that the rate constants for activating the first burst are slower than that for activating the subsequent bursts for genes B and C, which suggests the necessity of using three-state models for describing both genes (**Supplementary Fig. 5A**).

Besides state transition rate constants, additional parameters are needed to describe the circuit dynamics. For example, we needed to estimate RNA synthesis rate during the ON state. Because each burst produces an integer count of pre-mRNA molecules, we estimated each molecule’s fluorescence intensity by performing fast Fourier transforms on the density function of the bust amplitude (**Supplementary Fig. 5F**), allowing us to approximately estimate RNA synthesis rate (Methods).

Together, based on the experimental time-lapse data, we parameterized three-state models for describing transcriptional bursting dynamics of each gene in the synthetic three-gene circuit, allowing us to investigate further what parameters affect the propagation of dynamic signals in the cascade.

### Dissecting temporal gene-gene interactions using model simulations

Using the model, we performed stochastic simulations to study the temporal transcriptional dynamics of the circuit in response to a step signal analogous to the experiment (Methods). We successfully recapitulated key features of the experimental data, including the sequential activation of transcriptional bursts in the cascade (**Supplementary Fig. 6A-B**), and exponentially or peaked response time and dwell-time distributions (**Supplementary Fig. 6C**), which are unaffected when doubling the value of *K_T_* parameter (**Supplementary Fig. 6D**). The simulated results also captured the propagation of the first, but not subsequent, burst of transcription from the upstream gene to its immediate downstream target (**Supplementary Fig. 6E** top).

By examining the simulated temporal traces of molecular species in the circuit, including mRNA and protein (**Supplementary Fig. 6B**), we sought to identify parameters influencing the transmission of dynamic bursts from the upstream regulator to downstream target. Focusing on the interaction between A and B, we speculated that the transmission of burst from A to B occurs only during the transient stage of the circuit’s response to the external signal, or when the protein level of gene A has not yet saturated gene B’s dose response curve. In contrast, when protein A level saturates, additional bursts of gene A’s transcription would not alter the kinetics of gene B’s bursting. To test this hypothesis *in silico*, we performed the simulation by lowering mRNA and protein stabilities of gene A (Methods), and observed an overall enhanced propagation of bursts (**Supplementary Fig. 6E-F**). The result indicates that the propagation of transcriptional burst from the upstream regulator to downstream target is enhanced when the dynamics is more non-stationary.

While simulations provided insights into the propagation of dynamic transcriptional bursts in this linear cascade, our model cannot capture the observation that gene A appears to be switched off at later time points in some cells even in the presence of doxycycline (**Fig. 3C** and **Supplementary Fig. 3B**), indicating that mechanisms in addition to transcription factor binding likely control the activity state of the gene.

### Interrogating temporal interactions in the presence of combinatorial regulation

We next sought to construct and analyze a different synthetic circuit to interrogate how transcription factors temporally interact in the presence of hierarchical organization and combinatorial regulation, i.e., the two features of human gene regulatory network^54^.

We constructed a monoclonal U2OS cell line containing a five-gene circuit constituting a branched regulatory cascade (**Fig. 4A**). This circuit can respond to an external signal (i.e., doxycycline), and the last gene is combinatorially regulated by two upstream regulators, which acts as a three-input AND gate (two regulators plus rapamycin). Using this circuit, we can study how dynamic regulatory signals are propagated along two parallel branches and whether logic gate shapes burst propagation. With five distinct barcodes (**Fig. 4A**), we were able to resolve five genes in single cells (**Supplementary Fig. 7A**), and quantify the transcriptional dynamics of each gene in response to a step increase in the external signal (**Fig. 4B-C**, **Supplementary Video 4**). Population-averaged trajectories (**Fig. 4D**) revealed sequential and cascading activation over a time course of >1.5 days. Interestingly, the two parallel branches appeared to be activated at different rates (i.e., gene B and gene E were not synchronously activated, see **Fig. 4D**).

**Figure 4.**
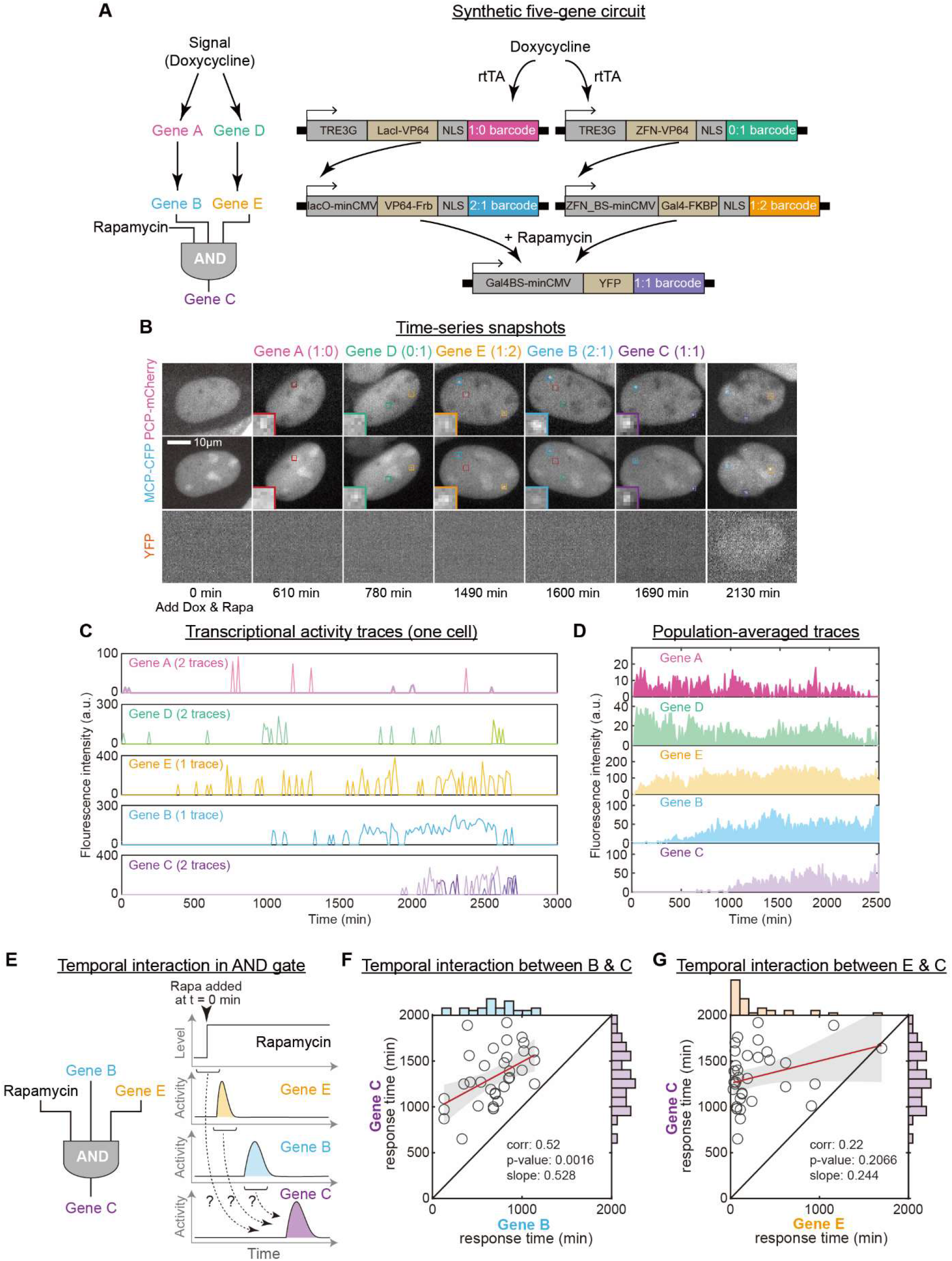
Multiplexed transcriptional imaging reveals imbalanced transcriptional burst propagation in a synthetic five-gene circuit. **(A)** Schematic of a synthetic five-gene circuit. Doxycycline induces the expression of LacI-VP64 (gene A) and ZFN-VP64 (gene D), which then activates VP64-FRB (gene B) and Gal4-FKPB (gene E). In the presence of rapamycin, the last two proteins together activate YFP (gene C). **(B-C)** Time-lapse images (B) and corresponding transcriptional activity traces (C) of a representative single cell containing the five-gene circuit. Doxycycline (1 μg/ml) and rapamycin (100 nM) were added at t = 0 min. See also **Supplementary Fig. 7A** and **Supplementary Video 4**. **(D)** Population-averaged transcriptional activity traces of individual genes in the circuit (n = 33 cells). Same experimental condition as in (B). **(E)** Illustration of temporal gene-gene interactions through the AND gate at the level of individual transcriptional bursts. **(F-G)** Evidence for imbalanced transcriptional burst propagation through the AND gate. Same experimental condition as in (B). Scatter plots of response times for two gene pairs are shown (n = 45 events for each panel). Response time refers to the start time of the first transcriptional burst after dox addition (see **Fig. 3G** for schematic). Red line is the least-squares fit of the data and gray shadow denotes 95% CI of the fitted line. The target, gene C, displays a significant correlation with upstream regulator gene B (panel F), but not with gene E (panel G), indicating that only the transcriptional burst of gene B propagated to gene C. See also **Supplementary Fig. 7B-D**.

To quantify signal propagation through the AND gate (**Fig. 4E**), we performed correlation analysis analogous to the three-gene circuit (**Supplementary Fig. 4B**) between the two upstream regulators (gene B and gene E) and their co-regulated target gene (gene C). Note that when rapamycin is present throughout the time course, the three-input AND gate effectively becomes a two-input AND gate. We found that the target gene C’s first burst exhibited a significant correlation with its upstream regulator gene B (**Fig. 4F**), but not with its other regulator gene E (**Fig. 4G**). Because target gene C can only be turned on when these two regulators are both present (**Fig. 4A**), such an imbalanced dynamic transmission is likely due to the slower arrival of gene B’s first burst compared to that of gene E (**Fig. 4C-D**), suggesting that the slowest-expressing regulator in this AND gate determines the initial burst timing of the target gene.

To test this picture, we sought to artificially control the timing of the rapamycin signal, which is an input signal of the AND gate and can be externally manipulated (**Fig. 4A**). Because the arrival time of the target gene C’s first burst is between 600~2000 min in the preceding experiment (**Fig. 4F-G**), we hypothesized that administering rapamycin signal within this time window would set a lower temporal bound for target gene’s response (**Supplementary Fig. 7B**). As expected, we found that adding rapamycin at ~1000 min, instead of adding rapamycin at time zero in the preceding experiment, caused the target gene C to delay its first burst until after the rapamycin signal (**Supplementary Fig. 7C**). Importantly, gene C still exhibited a significant correlation with its regulator gene B (**Supplementary Fig. 7C**), but not with its other regulator gene E (**Supplementary Fig. 7D**), indicating the robustness of imbalanced burst propagation to the perturbation of the rapamycin signal.

Together, these results demonstrated that the propagation of bursts of upstream regulators to their downstream targets may depend on other regulatory inputs controlling the target, and that a regulatory relationship between two genes may not necessarily result in a correlational relationship between them.

## DISCUSSION

Gene regulatory circuits are inherently dynamic, yet there have been few tools for temporal analysis of complex gene circuits in single cells, partially due to the limited number of distinguishable fluorescent proteins. In this work, we developed a multiplexed transcriptional imaging method based on ratiometric RNA labeling using PP7 and MS2 stem-loops. We showcased the capability of this method by interrogating multi-gene interactions in synthetic circuits that recapitulate prominent features of human gene regulatory network^54^ such as hierarchical organization and combinatorial regulation.

Compared to analyzing genetic circuits at the level of protein products by using fluorescent proteins^23–25^, analyzing circuits at the level of transcriptional activities provides several advantages. Importantly, transcriptional activity-level analysis avoids influences from post-transcriptional regulation, yielding direct information regarding gene regulation. Compared to other tools for RNA imaging^51,63,64^, although RNA stem-loop-based approach requires relatively more complicated cloning and genetic manipulation procedures, it allows multiplexed barcoding and quantitative reporting of nascent transcriptional activities. With this method, one could examine multi-gene circuits in the same single cells at a high temporal and spatial resolution.

While temporal gene-gene interactions can also be studied with fluorescent protein-based reporters^29,65^, the current approach describes multi-gene interactions at the level of nascent transcriptional activity bursts. With such a temporal resolution, we revealed that gene-gene interactions can be best captured during the non-stationary phase of the system (i.e., when the dynamics have not yet reached a steady state) **(Fig. 3F-G**). This finding suggests that it is critical to consider the measurement timing when inferring gene regulatory relationships using static single-cell snapshots (such as single-cell RNA-seq data). More specifically, considering the linear gene regulatory cascade in **Fig. 3A**, one could not obtain a high correlation between the regulator gene and the target gene when their mRNA (or protein) levels in individual cells are measured at later time points of the time course (**Supplementary Fig. 8** and Methods). Additionally, gene-gene interaction can also be influenced by combinatorial regulatory logic (**Fig. 4**). Altogether, we illustrated several scenarios where a casual regulatory relationship might not necessarily confer a correlational behavior, highlighting potential challenges for inferring gene regulatory relationships using static single-cell gene expression data^17,53^.

We believe that the presented method should be highly scalable by incorporating additional RNA binding proteins^66^, other RNA labeling techniques^64^, or new multiplexing modalities. While it currently utilizes two channels to analyze five genes simultaneously, further developments towards expanded multiplexing capacity may allow one to temporally decode large natural or synthetic regulatory networks in live single cells.

## MATERIALS AND METHODS

### Plasmid construction

All PP7/MS2 barcode cassettes were constructed from two building blocks, i.e., 2xPP7 stem-loops and 2xMS2 stem-loops, which were based on the literature ^49^ and were synthesized. Due to the repeated nature of the sequences, cassettes were cloned based on isocaudomers digestion and T4 ligase ligation (NEB), and were checked by Sanger sequencing. Other plasmid components (see Supplementary Table 1) were synthesized or PCR amplified with PrimerSTAR MAX DNA polymerase (TAKARA) and checked by Sanger sequencing. Backbones were linearized with restriction endonucleases (NEB). All the DNA fragments were cloned into plasmids by Gibson assembly or by ligation using T4 ligase (NEB). Plasmids were replicated in DH5α cells (CWBIO) using standard protocols. Transgenes were cloned into the plasmids from the PiggyBac transposon system, except PCP-3×mCherry and MCP-3×CFP, which were cloned into the plasmids from the lentiviral packaging system (a gift from P. Wei). DNA sequences of barcodes and synthetic components used in our synthetic circuits are listed in Supplementary Table 1.

### Cell transfection and viral transduction

Transgenes were integrated using standard transfection or electroporation protocols, except for PCP-3×mCherry and MCP-3×CFP, which were integrated into U2OS cells through viral transduction. For viral transduction, lentiviral particles were produced by transient transfection of HEK-293T cells using polyethylenimine MW 25 000 (PEI, Polyscienc). U2OS cells were then transduced with lentiviral particles and were harvested 2 days post-transduction. Stable cell lines were constructed and utilized for subsequent integrations of synthetic genes carrying barcode cassettes. For transient transfections or stable integrations of single barcodes into U2OS cells, plasmids were transfected using PEI. Transfections were performed in 12-well plates, where cells reached ~80% confluency at the day of transfection. For transfection of one well, 2 μg DNA and 6 μl of 10 mM PEI solution were used. For stably integrating dual barcodes into cells, plasmids were transfected using Lipofectamine LTX with PLUS™ (Invitrogen). Transfections were performed in 12-well plates with ~80% cell confluency at transfection. 2 μg DNA, 2 μl Plus reagent, 6.0 μl LTX reagent were used for each well. For integrating three- or five-gene synthetic circuits into cells, plasmids were transfected into cells via electroporation following manufacturer’s cell type-specific protocol (NAPA21 TypeII Electroporator, NAPA GENE). 1×10^6^ U2OS cells were electroporated with 20 μg (three-gene circuit) or 40 μg (five-gene circuit) plasmids for each reaction. Cell culture conditions were as described ^44^.

### Stable cell line construction

Stable cell lines were obtained after antibiotic resistance selection and fluorescence-activated cell sorting (FACS), except that monoclonal cell lines were obtained from single cells deposited into 96-well plates using FACS. Prior to sorting, the inducible fluorescent protein was induced by adding 1 μg/ml doxycycline (Clontech) alone or together with 100 nM rapamycin (Harveybio) (for five-gene synthetic circuits) into the culture media for 12 hours (for cell lines with one or two barcodes) or for > 24 hours (for cell lines with synthetic circuits). Note that a longer time was needed to ensure the full activation of gene cascades. Cells with positive fluorescence signals were gated and deposited by FACS. The resulting polyclonal cells or single cells were cultured and expanded in antibiotic-containing and doxycycline-free culture media

### Cell lines used in figures

For Fig. 2A, five different polyclonal cell lines were used. Each cell line was integrated with a reporter gene fused with a distinct barcode (pTRE3G-Citrine-barcode), together with plasmids carrying rtTA for inducible expression of the barcoded reporter. For Fig. 2C and Supplementary Fig. 2B, polyclonal cell lines integrated with two types of reporter genes fused with distinct barcodes were used (pTRE3G-Citrine-2:1 and pTRE3G-iRFP-1:2, or pTRE3G-Citrine-0:1 and pTRE3G-iRFP-1:0). These cell lines were also integrated with plasmids carrying rtTA. Details of other cell lines have been appropriately described in various locations.

### Time-lapse fluorescence microscopy

Cells were seeded into wells of a 24-well glass-bottom plate (Eppendorf) ~12 hours prior to imaging, and reached ~60-70% confluency at the time of imaging. Time-lapse microscopy was performed on an automated inverted microscope (Nikon Ti-E) using a Plan Apo Lambda 40x objective. The microscope is equipped with a white-light LED (Lumencor SOLA), standard fluorescence filter sets (Chroma and Semrock), an automated sample stage, and a sCMOS camera (Hamamatsu ORCA-Flash4.0V2). The glass-bottom plate was maintained in an environmental chamber under 37°C and 5% CO2. Multi-color images were automatically acquired using Micro-Manager program^67^ at different frame rates for different types of experiments. For Single barcode experiments, the frame rates are 20 or 30 min/frame and for synthetic circuits the frame rates are 10 min/frame. Images were acquired continuously for ~10 hours (for single- and dual-barcode cell lines) or for 40 hours (for cell lines with synthetic circuits).

### Time-lapse image analysis

Fluorescence images were loaded into a custom MATLAB GUI and were analyzed semi-automatically using the graphic user interface created by the software. Each nascent transcription site was first identified manually, and then the software automatically identified the pixel location with maximum fluorescence intensity. The intensity of each transcription site was calculated by subtracting the signal intensity by the background intensity. For calculating the signal intensity, a 3×3-pixel square was defined around the local maximal pixel, and the intensities of the top four pixels were averaged. For calculating the background intensity, a 9×9-pixel square was defined around the local maximal pixel, and the pixels with intensity values ranking from 20% to 50% were regarded as background pixels whose intensities were averaged. Transcription site traces were manually marked for each frame and the resulting pixel locations were automatically tracked by the program. Each gene could have multiple transcription sites, corresponding to multiple integration events. Note that there are often scenarios where the transcription site did not appear on specific frames, and the intensity of the specific gene locus for these frames would be defined as zero.

Transcriptional burst is defined as a continuing series of transcription events. In our measurement, a burst starts from a frame that the intensity is larger than zero and the intensity of the last frame of the burst is zero or smaller than zero. That said, a burst ends at the first frame that the intensity is 0 or smaller than 0 after the initial frame. We can then define the burst duration and interval as the time between start and end frames, and between end and start frames, respectively.

### Estimation of exponential decay constant

The distributions of the response time of gene A, burst interval, and burst duration of all measured genes were exponential-like. As these reactions could be single-step, we assume those distributions are exponential and we could estimate the decay constants *λ* by moment estimators. Because our measurements were performed every 10 minutes, the event occurring within 10 minutes could not be detected. The decay constant *λ* of the exponential distribution is estimated by moment estimation: *λ* = 1/[E(*x*)-10], in which *x* is the time variable that obeys exponential distribution.

### Estimations of transcription rate constant

To construct the kinetic model of the three-gene circuit, we needed parameters describing the rate of mRNA transcription (*k_T_*) for each gene. Because parameter *k_T_* is not as important as other parameters and the choice of its value does not affect our conclusion, we here provide a rough estimation of *k_T_*. To do so, we needed to convert the fluorescent intensity of nascent transcription sites into mRNA molecules that are being produced.

Assuming that transcription occurs at a constant rate *k_T_*, the mRNA count detected within every 10-minute duration should obey Poisson distribution. Correspondingly, the distribution of fluorescent intensity should be unimodal and long-tailed. We found that the fluorescent intensity distributions are indeed unimodal and display some periodicity, indicating the presence of discrete counts of mRNA molecules with the intensity of each mRNA corresponding to the period. To extract the potential period of such distributions, we first estimated the density of the intensity by Kernel density estimation with two different bandwidths. For gene A the bandwidths are 2 and 20, and for genes B and C the bandwidths are 4 and 40. These bandwidths were chosen to better visualize the periodicity with the smaller bandwidth, and to better visualize the unimodal distribution with the larger bandwidth. We estimated the period by performing a fast Fourier transformation of the difference between these two densities. We ensured that most mRNA counts are larger than 0. After obtaining the density of mRNA counts, *k_T_* was estimated by maximum likelihood estimation.

### Gene B in the three-gene circuit

To describe the regulatory relationship between upstream regulator and downstream target gene, we first generated scatter plots using upstream transcription rate (i.e., mean upstream gene’s transcription site intensity before the response of downstream target) versus target’s response time (Supplementary Fig. 5D). We found that gene B’s response is slower when upstream transcription rate increases, which is counterintuitive. To explain this, we found that the dots in the scatter plot could be divided into two clusters, indicating two different activation kinetics for gene B loci. We then used mixture Gaussian model to fit (MATLAB function fitgmdist) the log(upstream transcription rate) versus response time data to generate two clusters. By doing so, we effectively divided gene B loci into two distinct groups. In each group, gene B responded faster when upstream transcription rate increases.

### Estimation of input functions

For model construction, we needed to estimate the input function that describes the dependence of *k_on1_* of the downstream gene on upstream TF concentration. Note that in contrast to *k_on1_*, which cannot be simply estimated by steady-state data, *k_on_* was estimated by the steady-state dwell time distribution. To estimate input function, we first estimated the TF concentration. Because we only measured transcriptional activity, we assumed that translation occurs at a constant rate *k_P_* of 10 protein per mRNA per min ^68^, and did not consider the degradation of both mRNA and protein. We then used the cumulative sum of transcribed mRNA counts as the mRNA counts at given time points, and the cumulative sum of these counts multiplied by *k_P_* was used as TF protein counts. The rationale for doing so is that we only needed an estimation of relative, rather than absolute, protein counts.

We next used a Hill equation to describe the input function:

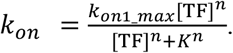

In this equation, *n* is the Hill coefficient, *k_on1_max_* represents the maximal rate of *k_on1_* when TF saturates, and *K* represents the dissociation constant. In our experiment, we induced the expression of the TF and measured the transcriptional dynamics of both the TF and the target gene. Because TF protein accumulates over time, the turning-on event of the target gene should be modeled as an inhomogeneous Poisson process, where *k_on1_* varies across the process. Thus, the probability of the first passage time *T_1_* (i.e., the appearance of the first transcriptional burst of the target gene or the response time) can be described as following:

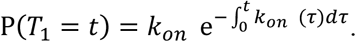

It is noted that when *k_on1_* is constant, the above equation becomes:

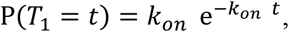

which describes a Poisson process and the mean of *T_1_* (i.e., E(*T_1_*)) equals the inverse of *k_on1_*. The TF protein count monotonically increases with time, and thus *k_on1_* would similarly increase with time, which can be illustrated by the increase in 1/E(*T_1_*) as TF protein (or TF transcription rate) increases. Because of the relatively small sample size, we used kernel density estimation to estimate the mean of *T_1_* at the same upstream transcription rate. We then used the preceding data to estimate the parameters *k_on1_max_* and *K* in the above Hill equation using maximum likelihood estimation. Note that we chose a Hill coefficient (*n*) of 2 in order to achieve a better performance of the estimation process.

### Model simulation

We simulated the kinetic model for the three-gene circuit using experimentally estimated or empirical parameters. Because individual reaction steps occur as Poisson processes, an accurate method for simulating such a stochastic system would be the Gillespie algorithm^69^. However, it is computationally heavy and we do not need such a high temporal resolution, as we experimentally sampled the system every 10 min. A tau-leaping version of the Gillespie algorithm would help to improve the speed of simulation but it is still slow. Thus, we used Markov process to simulate this process, which could be fast when sampling every 10 min. More specifically, the products of transcription and translation reactions for every 10 min were simulated by drawing from Poisson distributions. In every 10 min, TF protein counts would determine *k_on1_* and *k_on_*, then the probability for turning-on and turning-off events occurring within 10 min could be calculated as the dwell time distribution is exponential. To simplify the simulation, we assumed there is only one allele of each gene. The degradation rates of mRNA and protein are unknown in our experiments so we took 0.00116/min (half-life of about 10 h) as the mRNA degradation rate^70^, and 0.000566/min (median turnover of the human proteome) as the protein degradation rate^71^.

### Model simulation with different protein degradation rates

In Supplementary Fig. 6E-F, we aimed to study how protein degradation rate would affect burst propagation. To do so, we simulated the model of the three-gene circuit using two sets of parameters in order to compare different degradation rates. More specifically, the degradation rates of the mRNA and protein were multiplied by a factor *k*. To ensure that the reduced concentration of protein can still activate the downstream gene, the *K* in the input function was divided by *k*. *k* was set to 1 and 100 for the scenarios of slow and fast degradation, respectively.

### Using simulations to analyze time-dependent correlation between genes

In Supplementary Fig. 8, we simulated the three-gene circuit using preceding parameters (Supplementary Fig. 5) to obtain 1,000 single-cell traces of all molecular species (pre-mRNA, mRNA, and protein) for each gene. Using these single-cell traces, we computed Pearson correlation between regulator gene and target gene at different time points after the addition of the signal at time zero. The correlation was computed for all three molecular species. Note that for typical gene regulatory network inference algorithms based on single-cell RNA-seq data^72^, correlation (or a similar measure such as mutual information) is calculated using mRNA levels of regulator and target genes, which is analogous to the middle panels of Supplementary Fig. 8B.

## Data and code availability

The data and code used in this paper can be downloaded from the following link: https://github.com/IndigoMad/Multiplexed-transcriptional-reporter

## ACKNOWLEDGMENTS

Y.L. acknowledges the supports from National Natural Science Foundation of China (Grant # 31771425) and National Key R&D Program of China (Grant # 2020YFA0906900, 2018YFA0900703). M.B.E. acknowledges the supports from National Science Foundation (Grant # 1547056; Grant # EF-2021552 under subaward UWSC10142). MBE is a Howard Hughes Medical Institute Investigator. We thank the flow cytometry core at the National Center for Protein Sciences at Peking University and the Quantitative Imaging facility at the Center for Quantitative Biology at Peking University for equipment supports.

## AUTHOR CONTRIBUTIONS

Y.L. and M.B.E. conceived the concept of the reporter; S.X., K.L. and L.M. performed the research; J.Z. and S.Y. contributed new reagents/analytic tools; S.X. K.L. and Y.L. wrote the manuscript with inputs from M.B.E.

---

**Supplementary Figure 1.**
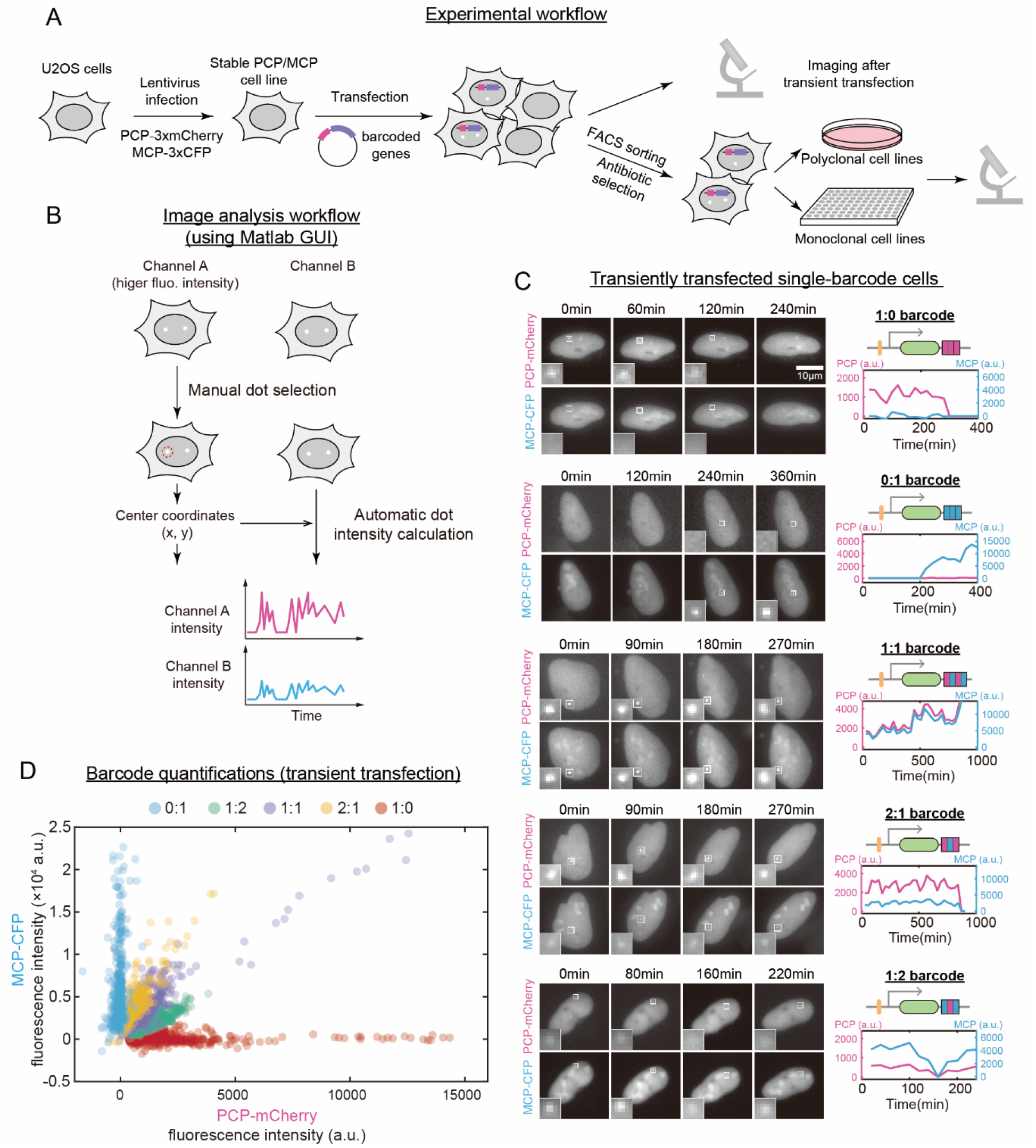
Construction and characterization of the multiplexed reporter system. **(A)** Flowchart illustrating experimental procedures. For experiments with transient transfection, cells were imaged ~24 hours after transfection. Stable cell line construction was accomplished by flow sorting and antibiotic selection. See Methods for details. **(B)** Flowchart illustrating image analysis procedures. Image analysis was performed using a custom Matlab graphical user interface (GUI). **(C)** Time-lapse images (left) and temporal traces (right) of representative single cells transiently transfected with a single type of genes fused with indicated barcodes. See **Fig. 1A** for barcode details. Doxycycline (1μg/mL) was added before t = 0 min. **(D)** Quantification of ratiometric barcodes using transiently transfected cells. U2OS cells transiently transfected with single type of barcode-labeled genes were analyzed (n = 15-20 cells for each barcode) and the fluorescence intensities for all barcodes were plotted together.

**Supplementary Figure 2.**
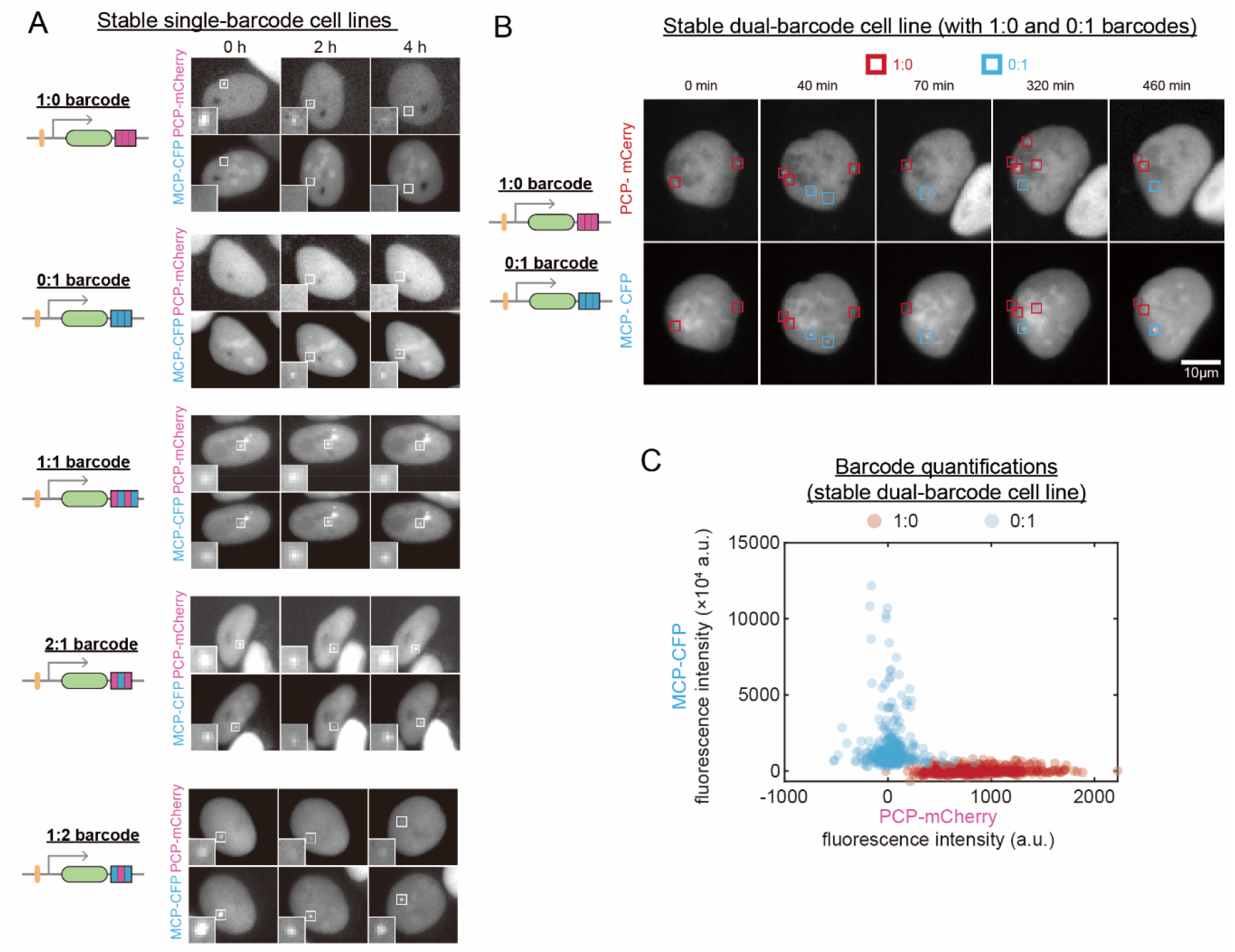
Transcriptional imaging of cells stably integrated with one or two barcode-labeled genes. **(A)** Snapshots of representative single cells stably integrated with genes fused with indicated barcodes. These cells are the same as in **Fig. 2A**. Doxycycline (1 μg/mL) was added before t = 0 min. See also **Supplementary Video 1**. **(B-C)** Two-color time-lapse images of a representative single cell integrated with two different genes carrying distinct barcodes (1:0 and 0:1 barcodes) (B), and the fluorescence intensities of barcoded gene loci (C, n = 11 cells). Doxycycline (1 μg/mL) was added before t = 0 min. See also **Supplementary Video 2**.

**Supplementary Figure 3.**
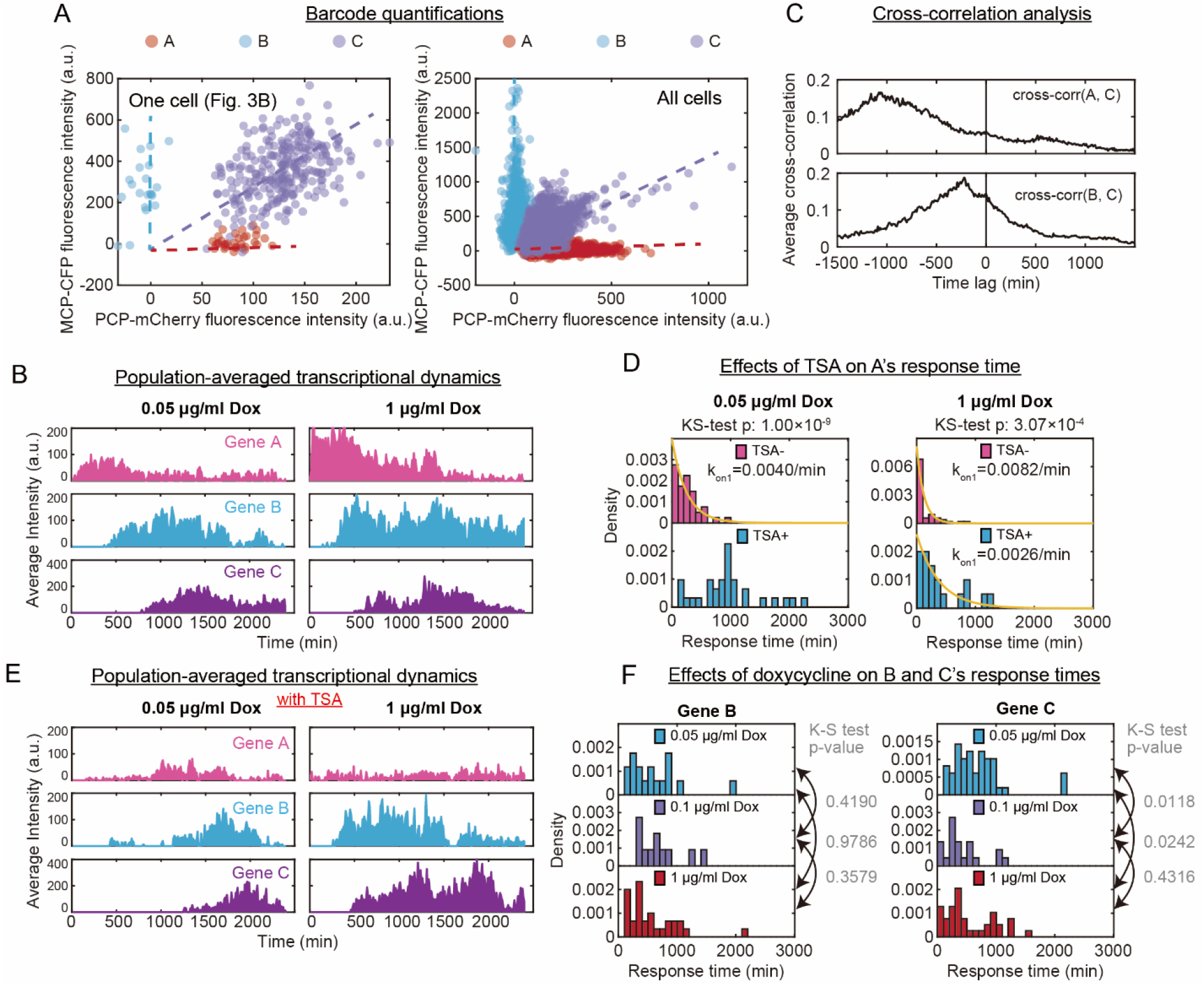
Characterizations of the three-gene circuit. **(A)** Quantification of ratiometric barcodes in cells stably integrated with the synthetic three-gene circuit. Barcode intensities of the single cell in **Fig. 3B** (left) and barcode intensities of all cells (right, n = 45 cells) are shown. Dashed lines are for visual guidance. **(B)** Population-averaged transcriptional activity traces of three genes under two indicated doxycycline conditions. **(C)** Averaged cross-correlation functions between gene A and gene C (top), and between gene B and gene C (bottom). Cross-correlation was calculated in individual cells and was averaged over 45 cells. **(D)** Distributions of the response times of gene A under indicated doxycycline conditions with or without TSA (100 nM). Note that TSA treatment at 0.05 μg/mL doxycycline altered the distribution from exponential to peaked distribution. n = 31 (left) and 20 (right) events for with TSA conditions. **(E)** Population-averaged transcriptional activity traces of three genes under two indicated doxycycline conditions in the presence of TSA. Note that TSA mainly affected the response time or activation level of gene A. **(F)** Distributions of the response time of gene B (left) and gene C (right) under indicated doxycycline conditions. n = 17, 11, 30 events for gene B, and n = 49, 22, 39 events for gene C.

**Supplementary Figure 4.**
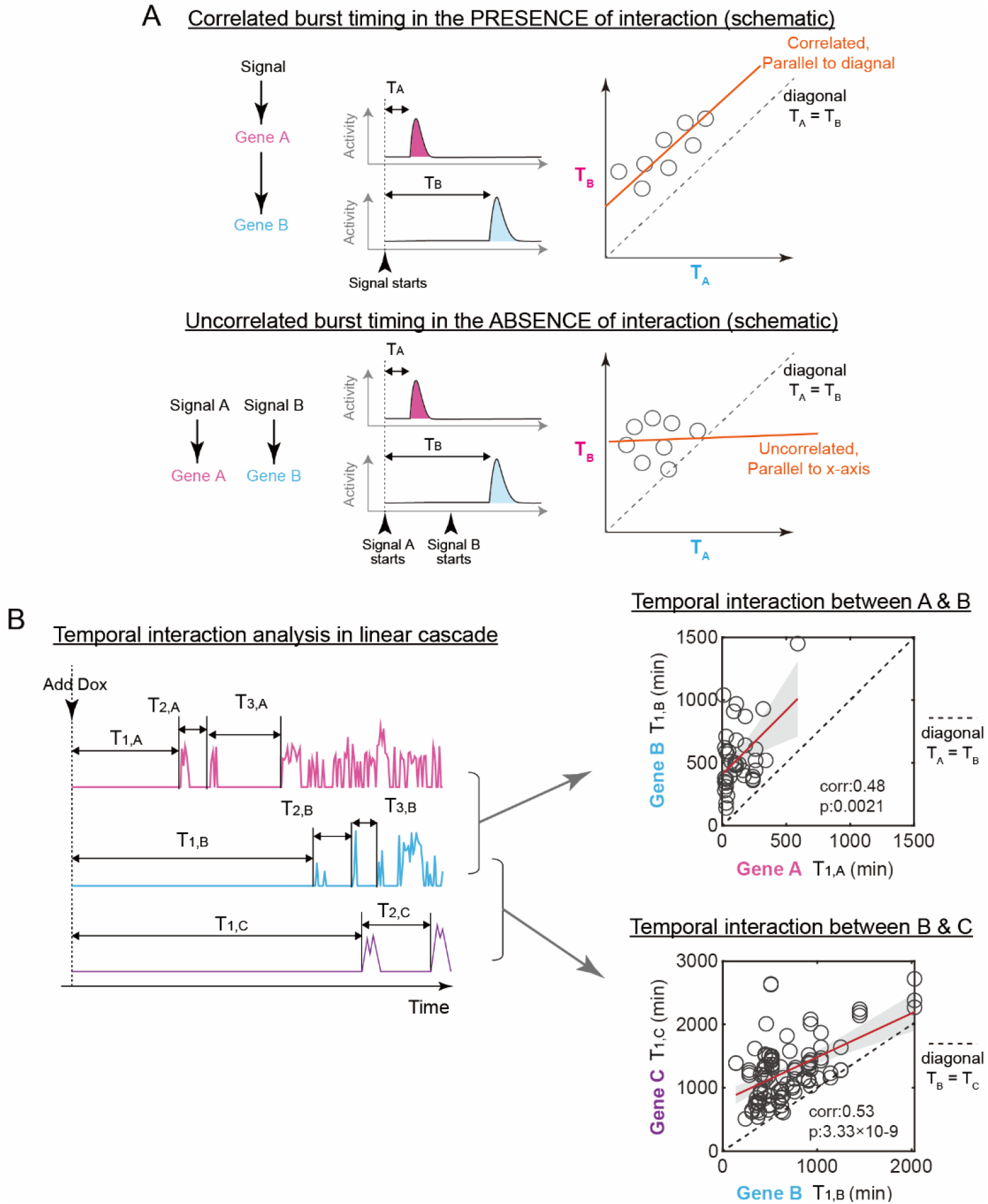
Analyzing temporal gene-gene interaction using transcriptional activity traces. **(A)** Schematics illustrating how the correlation between burst starting times can capture the presence or absence of regulatory interaction between two genes. In a circuit that a step-like signal activates gene A, and A regulates B, the initial transcriptional burst of gene A would occur before that of gene B, leading to correlated burst starting times of the two genes (top). In the absence of such regulatory interaction where gene A and gene B are independently activated by external signals, there would not be a correlated behavior (bottom). **(B)** Correlations between the starting times of first transcriptional bursts of upstream and downstream genes in the three-gene circuit. These two scatter plots correspond to the first two bars in **Fig. 3G**. n = 39 (top) and 110 (bottom) events. Red line is the least-squares fit of the data and gray shadow denotes 95% CI of the fitted line.

**Supplementary Figure 5.**
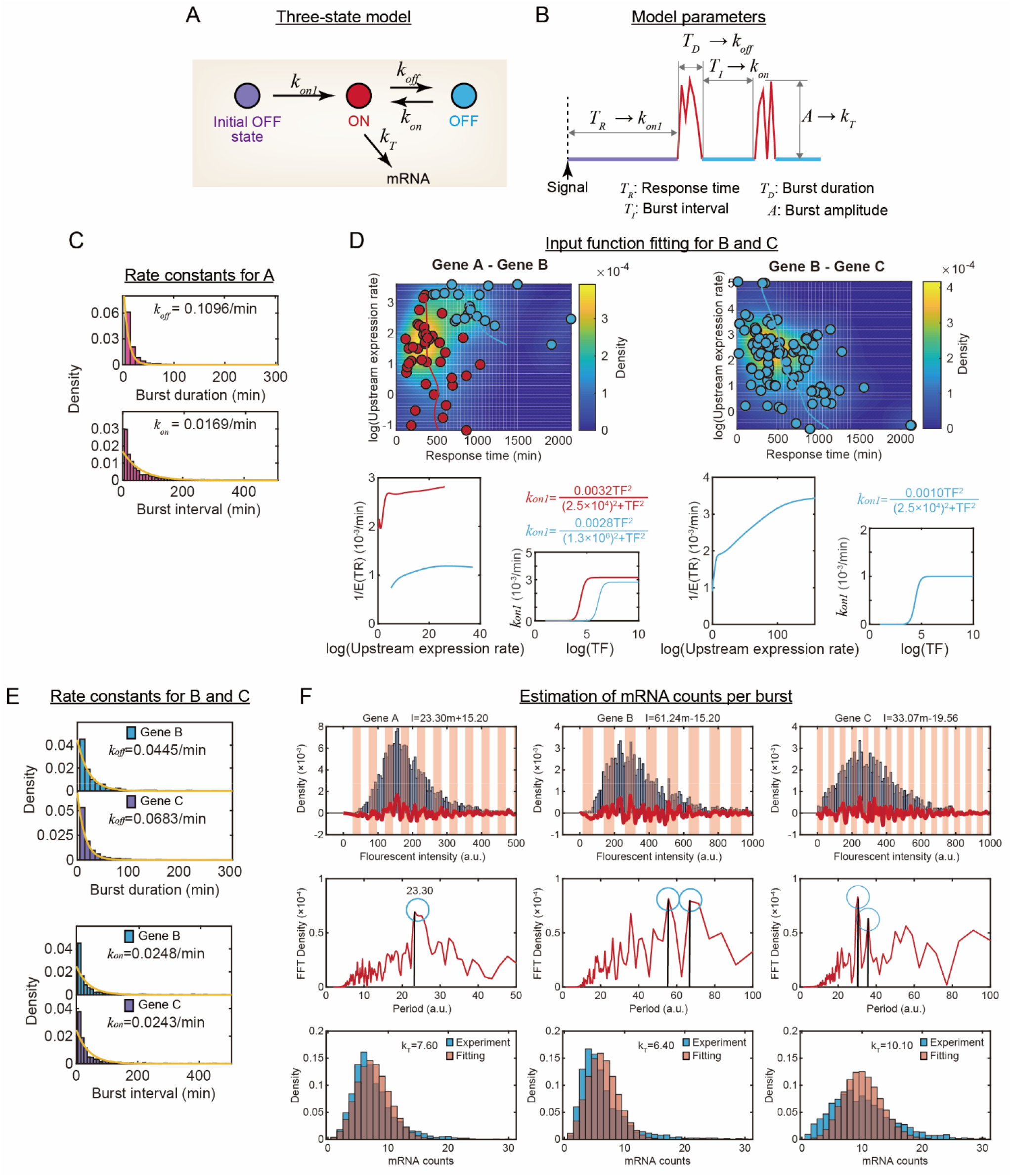
A three-state model to describe genes in the three-gene circuit and model parameter estimations using temporal data. **(A-B)** A three-state model and the schematic for model parameter estimations. In the model, there are two OFF states, including the initial OFF state before the addition of upstream signal and the steady-state OFF state (A). Upon the addition of the upstream signal, the gene turns on its first burst with a rate of *k_on1_*. During steady-state, it stochastically switches between ON and OFF states with indicated rates (*k_on_* and *k_off_*). When the gene is at the ON state, transcription occurs at a rate of *k_T_*. Using experimental traces, model parameters can be estimated as indicated (B). **(C)** Estimating gene A’s *k_off_* and *k_on_* parameters using burst duration (n = 1,426 events) and burst interval (n = 1,265 events) distributions. **(D)** Estimating promoter input functions for gene B (left) and gene C (right). Cells expressing gene B were divided into two subgroups based on the contour of upstream expression rate versus gene B’s response time (top left). In contrast, there is only one group for gene C (top right). For each group of cells, a Hill function was fitted to the data of upstream expression rate versus target gene’s response time, and the fitted input function was plotted (bottom). See Methods for details. **(E)** Estimating *k_off_* and *k_on_* parameters for B and C using burst duration and burst interval distributions. n = 565 (*k_off_*) and 489 (*k_on_*) events for gene B, and n = 853 (*k_off_*) and 741 (*k_on_*) events for gene C. **(F)** Converting fluorescence intensities into mRNA counts for the three genes in the circuit. For each gene, the period in the fluorescent intensity density was estimated and the fluorescent intensity was converted into mRNA counts. Top: the distribution of the fluorescence intensity (blue bars), the estimated density compared to local mean density (red line), and the intervals corresponding to one mRNA (red stripes). Middle: the period spectrum calculated by fast Fourier transformation, and the peaks of the density, i.e., the most possible periods (circles). In the presence of two peaks, the period was calculated as the mean of the peaks. Bottom: the distribution of the mRNA counts derived from the conversion of fluorescence intensities (blue) and the Poisson distribution (red) by fitting the mRNA count distribution. For gene A, the intensity of PCP-mCherry was used and for genes B and C, the intensity of MCP-CFP was used. Of note, the estimated *K_T_* parameter is likely not very accurate, albeit that this parameter is not very important for the dynamics (see **Supplementary Fig. 6C-D**).

**Supplementary Figure 6.**
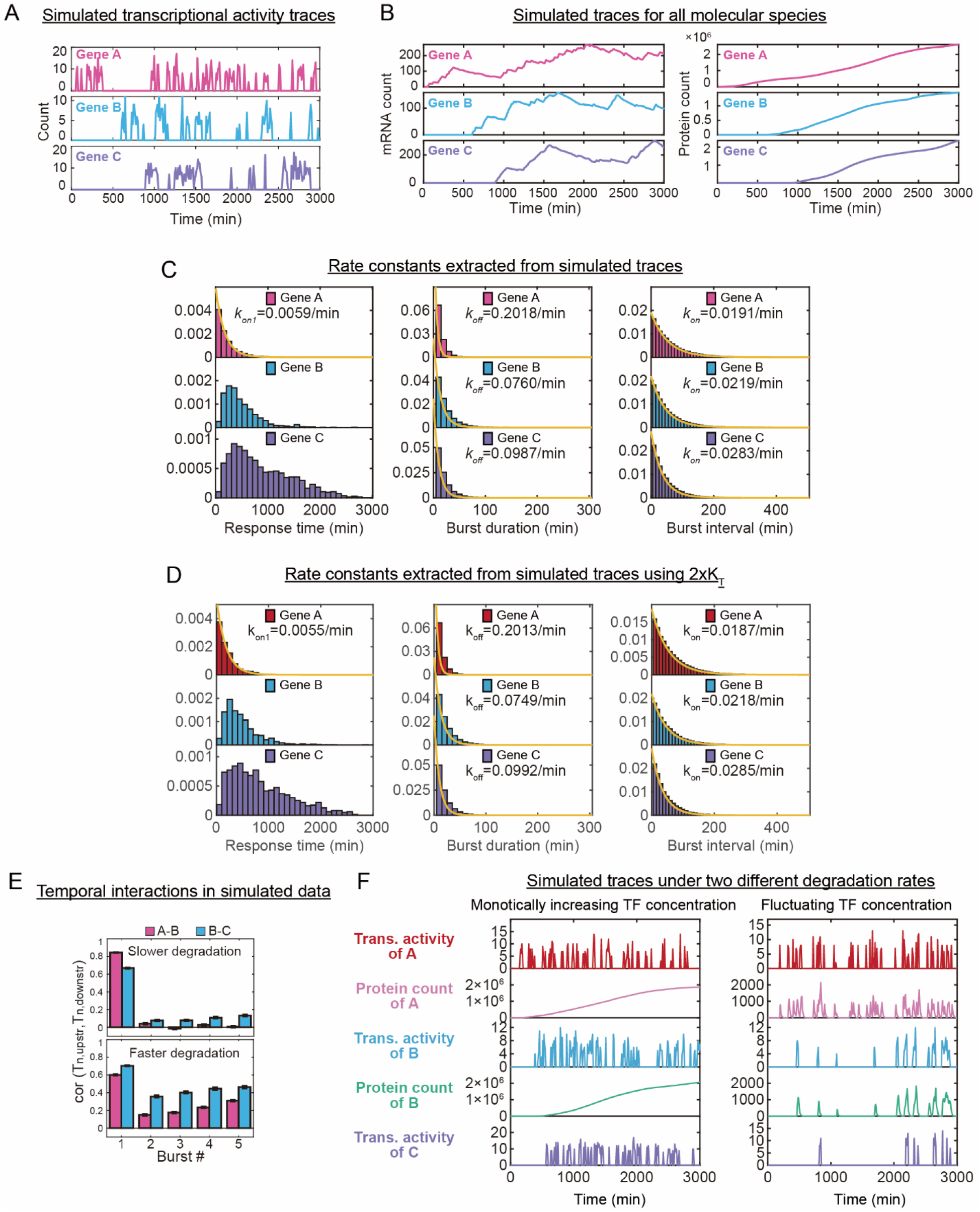
Dissecting temporal gene-gene interactions in the three-gene circuit using model simulations. **(A-B)** Simulated transcriptional activity (i.e., nascent mRNA count) traces (A) and mRNA and protein count traces (B) of the three-gene circuit using parameters that are mostly derived from experimental data. See also Methods. **(C)** Distributions of response times and dwell times of the simulated traces. From these distributions, parameters of the kinetic model can be quantified, and the resulting parameters are close to the input (experimental) parameters. **(D)** Analogous to (C) when parameter *K_T_* was set to 2×*K_T_* in the model. **(E-F)** Quantifications of burst propagation in two different groups of simulated traces (E). The two groups of traces were obtained by using two different parameter sets corresponding to different mRNA and protein degradation rates. Example simulated traces are shown in (F). See also Methods. Burst propagation was quantified analogous to **Fig. 3G**.

**Supplementary Figure 7.**
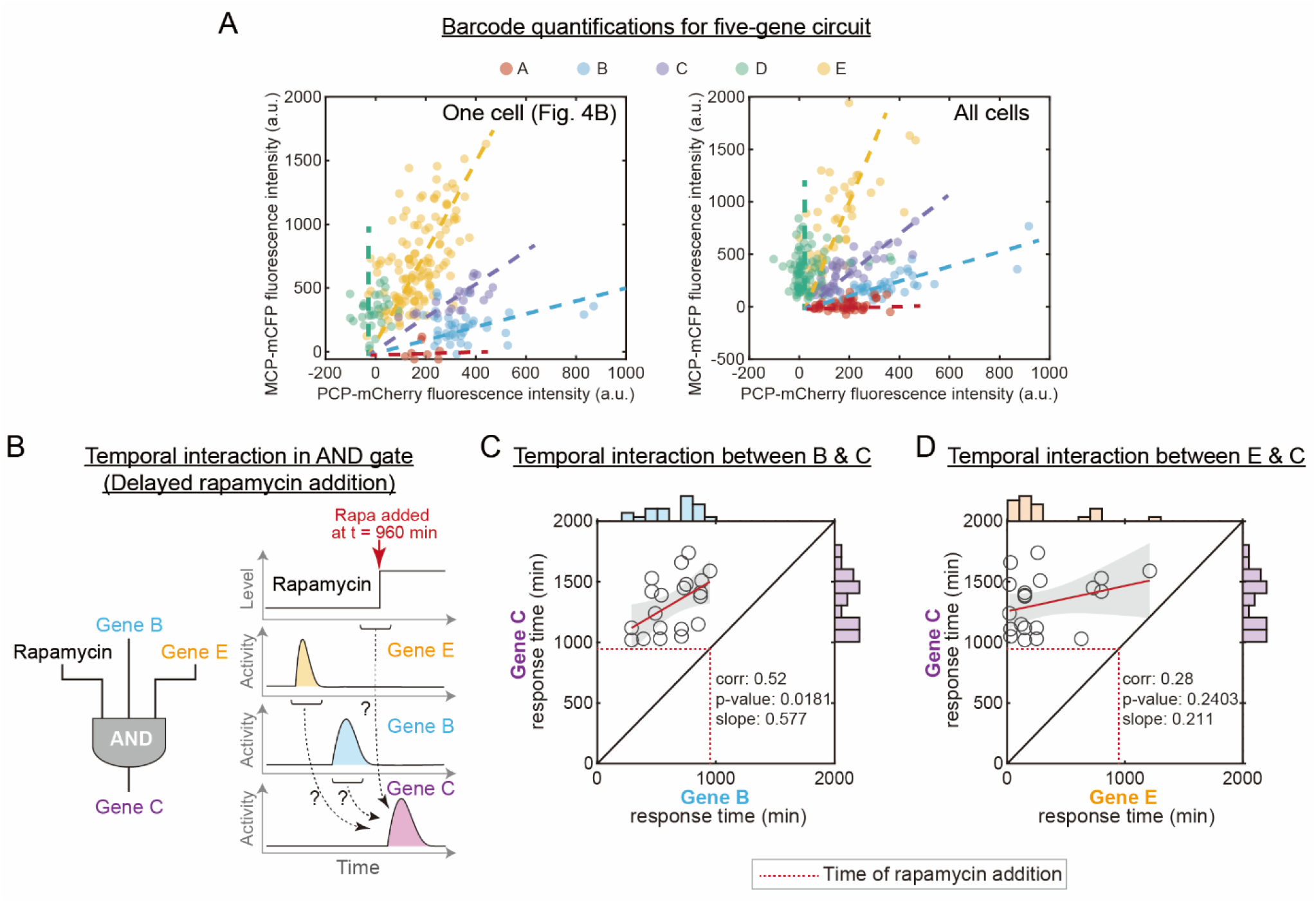
Barcode quantifications and additional analysis of temporal multi-gene interactions in the five-gene circuit. **(A)** Quantification of ratiometric barcodes in cells stably integrated with the synthetic five-gene circuit. Barcode intensities of the single cell in **Fig. 4B** (left) and barcode intensities of all cells (right; n = 33 cells) are shown. For the left plot, intensities from all time points are shown. For the right plot, only the data points with the highest intensities of each allele are shown for clarity. Dashed lines are for visual guidance. **(B)** Illustration of temporal gene-gene interactions through the AND gate at the level of individual transcriptional bursts. In this experiment, rapamycin was added at a later time point compared to the experiment in **Fig. 4E**. **(C-D)** Analysis of temporal interactions analogous to **Fig. 4F-G**. Rapamycin (100 nM) was added at t = 960 min (indicated by red dashed line). The target, gene C, displays a significant correlation with upstream regulator gene B (panel C), but not with gene E (panel D), indicating that only the transcriptional burst of gene B propagated to gene C. n = 37 (panel C) and 39 (panel D) events. It is noted that gene C could not be activated until after the addition of rapamycin (dashed red line).

**Supplementary Figure 8.**
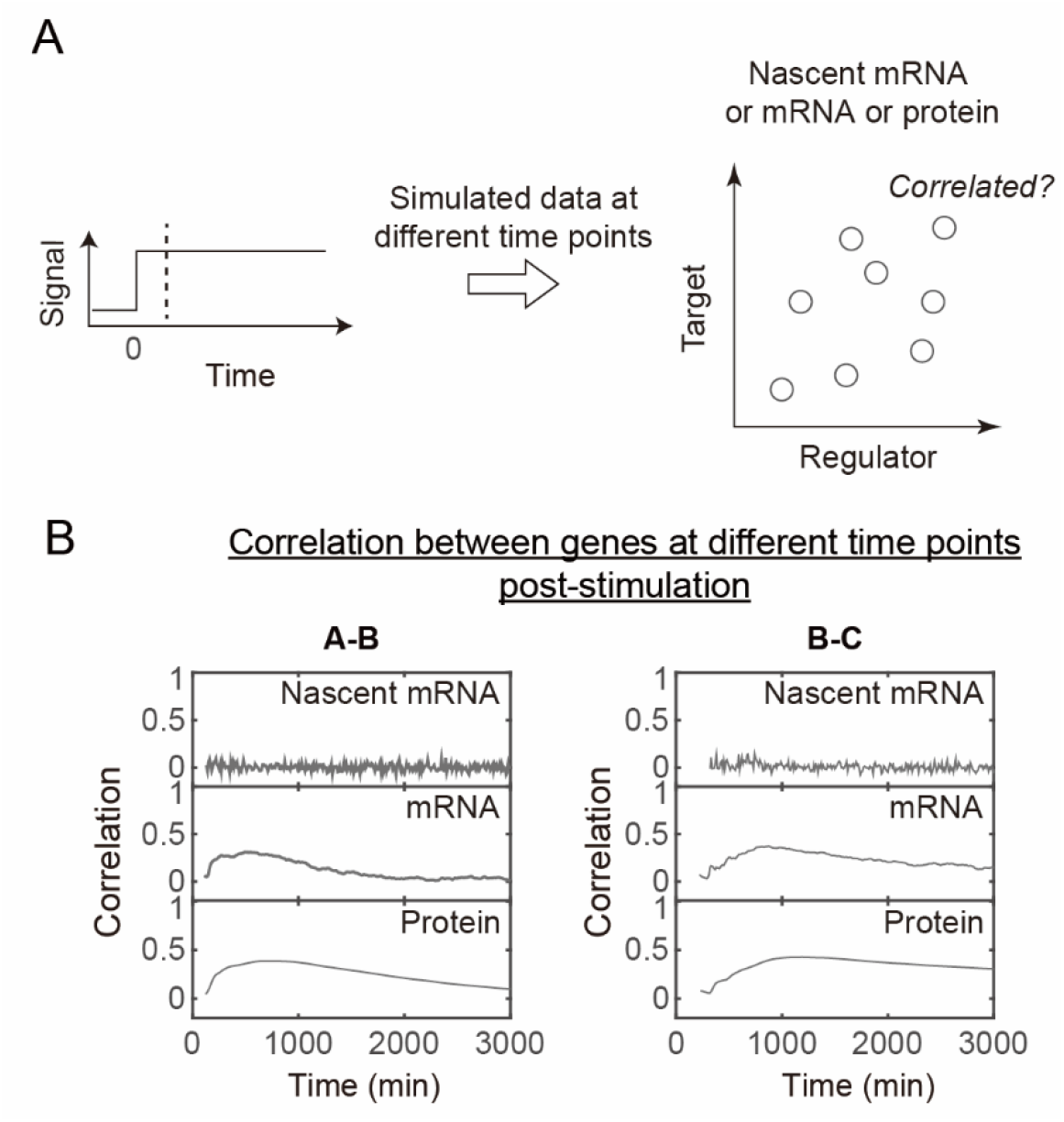
Correlation between genes is affected by the timing of the measurement. **(A)** Schematic illustrating how correlation is computed at different time points using simulated data. Dashed line on the left indicates data from a specific time point post stimulation by a step-like signal. Each dot in the scatter plot indicates data from a single cell. X- or Y- axis represents the level of pre-mRNA, mRNA, or protein of the indicated gene. **(B)** Correlation between regulator and target genes along time points post stimulation. The three-gene circuit was simulated as in **Supplementary Fig. 6** and traces analogous to **Supplementary Fig. 6A-B** were obtained for 1,000 cells. Pearson correlation coefficient was then computed across single cells at various time points post stimulation for each gene pair (A-B or B-C) at the level of pre-mRNA, mRNA, or protein. See also Methods. From these results, it is evident that at the level of mRNA or protein, the correlation between the regulator and its target is maximal during the transient phase post stimulation, which decreases as the system approaches the steady-state.

**Supplementary Video 1** | Representative two-color time-lapse movies of cells stably integrated with a single type of genes fused with five distinct barcodes. Doxycycline concentration is 1 μg/mL. Scale bars indicate 10 μm.

**Supplementary Video 2** | Representative two-color time-lapse movies and intensity traces of cells stably integrated with two types of genes fused with two indicated barcodes. Note that for clarity, we only plotted the fluorescence intensities of one transcription site per barcode. Doxycycline concentration is 1 μg/mL. Scale bars indicate 10 μm.

**Supplementary Video 3** | Representative two-color time-lapse movie and intensity traces of a cell carrying the three-gene circuit. Note that for clarity, we only plotted the fluorescence intensities of one transcription site per barcode. Doxycycline concentration is 1 μg/mL. Scale bar indicates 10 μm.

**Supplementary Video 4** | Representative two-color time-lapse movie and intensity traces of a cell carrying the five-gene circuit. Note that for clarity, we only plotted the fluorescence intensities of one transcription site per barcode. Doxycycline concentration is 1 μg/mL. Rapamycin concentration is 100 nM. Scale bar indicates 10 μm.

**Supplementary Table 1** | DNA sequences of barcodes and components used in our study.

